# Occasion setters determine responses of putative dopamine neurons to discriminative stimuli

**DOI:** 10.1101/799387

**Authors:** Luca Aquili, Eric M. Bowman, Robert Schmidt

## Abstract

Midbrain dopamine (DA) neurons are involved in the processing of rewards and reward-predicting stimuli, possibly analogous to reinforcement learning reward prediction errors. Here we studied the activity of putative DA neurons (n=41) recorded in the ventral tegmental area of rats (n=6) performing a behavioural task involving occasion setting. In this task an occasion setter (OS) indicated that the relationship between a discriminative stimulus (DS) and reinforcement is in effect, so that reinforcement of bar pressing occurred only after the OS (tone or houselight) was followed by the DS (houselight or tone). We found that responses of putative DA cells to the DS were enhanced when preceded by the OS, as were behavioural responses to obtain rewards. Surprisingly though, we did not find a population response of putative DA neurons to the OS, contrary to predictions of standard temporal-difference models of DA neurons. However, despite the absence of a population response, putative DA neurons exhibited a heterogeneous response on a single unit level, so that some units increased and others decreased their activity as a response to the OS. Similarly, putative non-DA cells did not respond to the DS on a population level, but with heterogeneous responses on a single unit level. The heterogeneity in the responses of putative DA cells may reflect how DA neurons encode context and point to local differences in DA signalling.

## 1. Introduction

Midbrain dopamine (DA) neurons are involved in the processing of rewards and reward-predicting stimuli. Their function of striatal DA has been proposed to be analogous to reinforcement learning reward prediction errors (RPEs), i.e. the difference between actual and predicted rewards (Eshel, Tian, Bukwich, and Uchida, 2016; Montague, Dayan, and Sejnowski, 1996; Schultz, 1998; Schultz, Apicella, and Ljungberg, 1993). The similarity of DA cell activity and RPEs has been demonstrated in simple classical conditioning tasks. Before learning, DA cells respond to the unconditioned stimulus, but not to the conditioned stimulus. However, during learning the response to the unconditioned stimulus gradually decreases, while the response to the conditioned stimulus gradually increases (Pan, Schmidt, Wickens, and Hyland, 2005). After many repetitions the response to the unconditioned stimulus may even cease, leaving only a response to the conditioned stimulus. This shift in the response from the unconditioned to the conditioned stimulus strongly resembles RPEs in simulations of classical conditioning using temporal-difference learning (Schultz, Dayan, and Montague, 1997), which has lead to the proposal that DA drives reinforcement learning in the brain (Schultz, 2016; Steinberg, Keiflin, Boivin, Witten, Deisseroth, and Janak, 2013).

While studies of DA responses to reward-predicting stimuli have used varying reward probabilities (Fiorillo, Tobler, and Schultz, 2003; Morris, Nevet, Arkadir, Vaadia, and Bergman, 2006) and magnitudes (Tobler, Fiorillo, and Schultz, 2005), reward-predicting stimuli in configural learning have received less attention (but see Waelti, Dickinson, and Schultz, 2001). For example, occasion setting, a phenomenon that has long been studied in the fields of learning and behavior (Fraser and Holland, 2019), has not been investigated in the context of DA neurons yet. In occasion setting a background stimulus (the occasion setter, OS) indicates that the relationship between a second stimulus (e.g. a discriminative stimulus, DS) and reinforcement is in effect. Thus, the presence of the OS indicates that reinforcement is possible, and the absence of the OS indicates that no reinforcement will occur. At a psychological level, the OS acts as a modulator of conditioned behaviour triggered by the second stimulus (Bonardi, Robinson, and Jennings, 2017; Trask, Thrailkill, and Bouton, 2017). Furthermore, the OS can facilitate reward-seeking, rather than merely simplifying ambiguous cue-reward pairings (Fraser and Janak, 2019).

How DA cells respond to the OS and DS in such tasks is interesting because standard temporal-difference learning models would treat the OS simply as the earliest reward-predicting stimulus in the sequence of events in a trial (Pan et al., 2005; Schultz et al., 1997), but neglect the importance of the combination of the OS with the DS. Therefore DA cell responses in occasion setting tasks may provide guidance for the development of more elaborate state representations in reinforcement learning algorithms that are employed by the brain (Russek, Momennejad, Botvinick, Gershman, and Daw, 2017). Furthermore, studying DA cell activity in occasion setting allows us to address whether DA cell activity also exhibits properties of a motivational signal. Recent evidence supported that slow, ramping increases in striatal DA are not due to corresponding firing rate changes in DA neurons (Mohebi, Pettibone, Hamid, Wong, Vinson, Patriarchi, Tian, Kennedy, and Berke, 2019). However, it has been noted that the type of behavioural task is a relevant factor for the expression of motivation signals (Berke, 2018), and they may not be as pronounced in head-fixed animals performing classical conditioning tasks compared to freely moving animals in operant tasks. As our OS task involved longer time scales, configural stimuli and operant behaviour, it was also suitable to test whether DA cell exhibits ramping firing rate increases as expected for motivational signals.

To study how DA cells take into account context for the processing of reward-predicting stimuli, we employed a behavioural task that involves occasion setting. Based on the RPE framework we predicted that DA neurons would respond to the OS, since it is the earliest stimulus in the chain of events leading to reinforcement. We also examined whether responses to a DS were gated by the presence or absence of an OS, and whether DA cells showed slow, ramping increases in firing rate towards the reward.

## 2. Methods

### 2.1. Subjects

16 Lister Hooded adult male rats (Harlan, UK) were housed in pairs on a light 12h: dark 12 h cycle and weighed 340 to 548 g when training began. Rats were allowed to consume water from 16.00 h to 17.00 h each weekday and from Friday 16.00 h to Sunday afternoon during experimental training. During this period, the rats’ body weights were monitored so that none fell below 85% of their free drinking weight, and all rats gained weight during the course of testing after minor losses due to surgery and the beginning of the regime of restricted water access. Following electrode implantation, rats were housed in isolation. All procedures conformed to the United Kingdom 1986 Animals (Scientific Procedures) Act, and ethical clearance was given by the Animal Welfare and Ethics Committee (AWEC) at the University of St Andrews.

### 2.2. Behaviour

In general, both the behavioural and neurophysiological methods reported here are similar to Wilson & Bowman (2006). Training and testing of rats occurred in sound-attenuated chambers (34 cm ⋅ 29 cm ⋅ 25 cm; Medical Associates Inc., St Albans, VT, USA), fitted with a video camera (Santec SmartVision, modelVCA 5156; Sanyo Video Vertrieb GmbH Co., Ahrensburg, Germany) for monitoring the rats’ behaviour. Each chamber contained a retractable lever, drinking spigot, houselight and piezoelectric buzzer (model EW-233A, Medical Associates Inc.). A reward magazine light (RL) was located in the interior of a reward magazine and consisted of a white LED (~2072 mcd luminosity). Sodium saccharin solution (0.25% w/v) was pumped out of the drinking spigot at 0.05 mL/s from a 50-mL glass syringe (Rocket, London) by computer-controlled syringe pumps (model PHM-100; Medical Associates Inc.)

### 2.3. Neurophysiology

The electrode arrays contained a movable bundle of four 50-µm stainless steel microwires coated in Teflon (impedance 0.4-1.3 MΩ). The microwires could be advanced by ~317.5 µm/turn in each recording session by turning an 80-thread/inch set screw (Small Parts Inc., Miami Lakes, FL, USA). The arrays weighed between 1.3 and 1.4g and measured 6mm along the mediolateral axis and 11mm along the anteroposterior axis. During recording sessions, the rat was connected to a preamplifier headstage using field effect transistors (input impedance 100 MΩ, unity gain voltage followers) which was in turn attached via a flexible cable to an electrical commuter (Stoelting Co., Wood Dale, IL, USA).

In order to remove noise, and lickometer artefacts, neuronal activity was recorded differentially from each of two pairs of wires. A custom built lickometer was also used to minimise lickometer artefacts (Malcolm McCandless, University of St Andrews) using a detection signal of sufficiently high frequency (> 5 kHz) that it could be filtered out. Amplification by 100 000x was obtained from each pair of wires using a Neurolog System (Digitimer Research Instrumentation) and frequencies <1 kHz and >5 kHz were attenuated by filters. Two Quest Scientific ‘Hum Bug’ digital filters (Digitimer) were used to eliminate 50 Hz noise. The differential activity from the two pairs of wires was finally digitised by the CED 1401+ data acquisition system using the associated Spike2 software (Cambridge Electronic Design, Cambridge, UK). Waveforms of putative action potentials were sampled at 20 kHz. Behavioural events were communicated from the MED-PC to the CED 1401+’s digital inputs for time-stamping. The temporal resolution of the MED-PC system was 2 msec.

### 2.4. Procedures

Rats were trained over a ~2 month period to reach the final stage of occasion setting training (see Figure 1) according to the stages described below. The initial training stages were adopted from a previous study (Wilson and Bowman, 2006). The later stages of training were modified from a previously published protocol (Holland, 1995). In this experiment, transfer of occasion setting properties was not examined as in the Holland paper.

**Figure 1.**
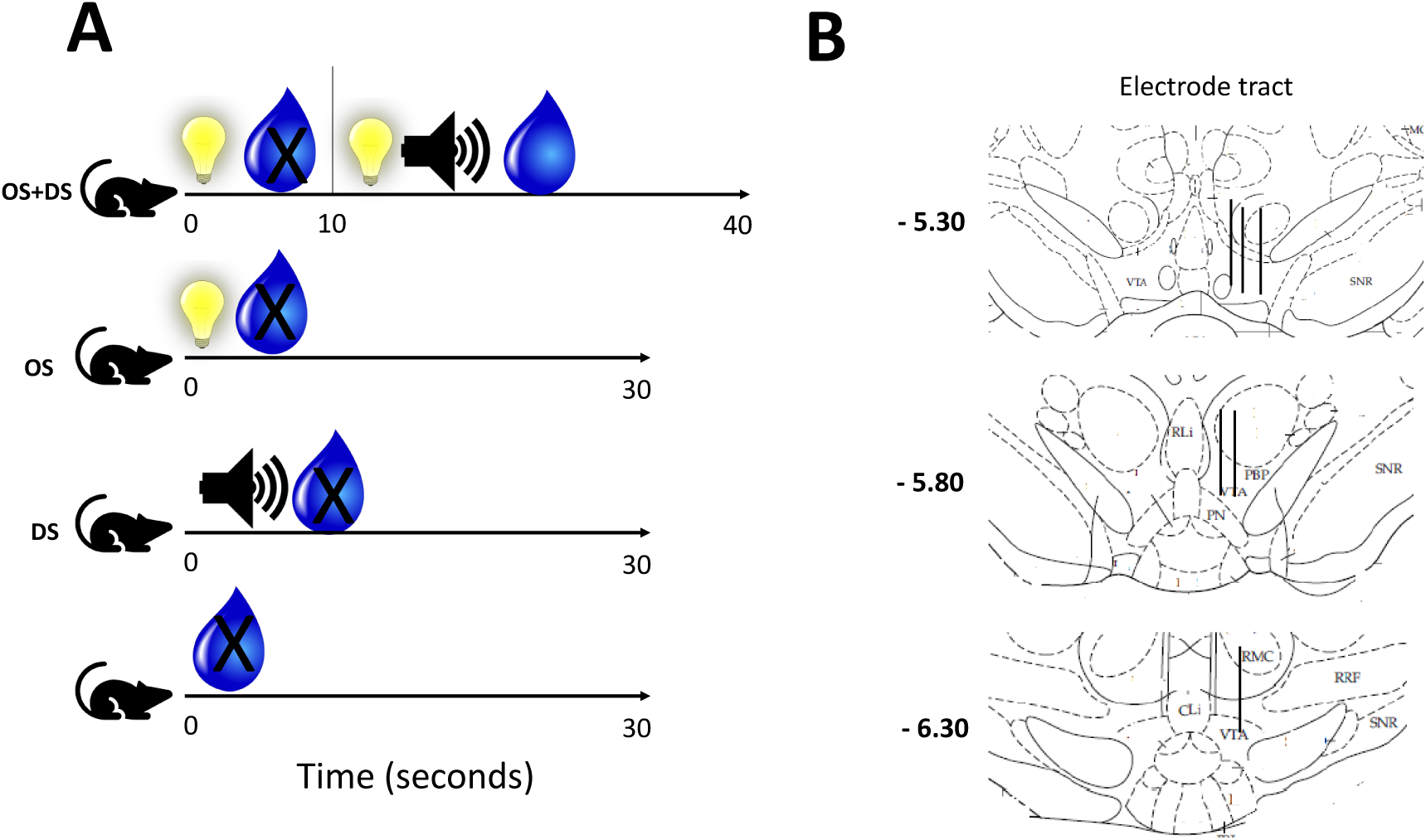
**A.** Top. Thirsty rats (n=16) were reinforced with saccharin solution for bar pressing only during OS (tone or light, counterbalanced) overlapping with a subsequent discriminative stimulus (DS, light or tone, counterbalanced) (i.e. between 10 seconds and 40 seconds of the trial). No reward was delivered if the animal bar pressed in the first 10 seconds of the trial (i.e. when only the OS was present). The rats could earn multiple reinforcers during periods in which OS and the DS were presented together. In all other trial types, bar pressing during OS alone (2^nd^ from the top), DS alone (2^nd^ from the bottom), or during no stimuli being presented (bottom), was not rewarded. B. Diagrammatic illustration of electrode tract position for the six rats during which putative dopaminergic and non-dopaminergic cells were recorded. Coronal sections are shown from −5.30mm to −6.30 with respect to Bregma.

**Figure 2:**
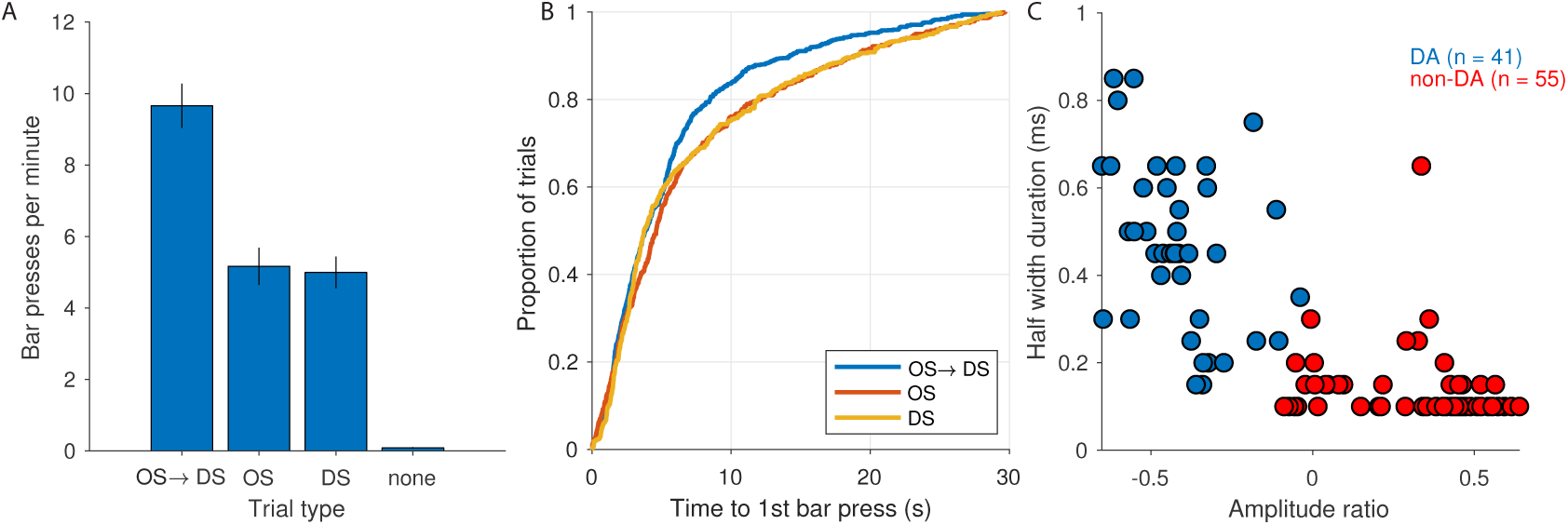
Behavioural data and cell type classification. (**A**) The mean number of bar presses per minute (±SEM) is shown for the different trial types (averaged across sessions and animals). For the trial type OS→DS the 30s period following the DS was considered; for the trial type OS the 30s period following the OS; for the DS trial type the 30s period following the DS; and for ‘none’ the 30s control period without any stimuli or rewards. (**B**) Response times are visualised as the cumulative distributions of the time between stimulus onset and the first bar press. For OS→DS trials this was the time between DS onset and bar press (same for DS trials, but for OS trials the response time was the time between OS onset and first bar press instead). Response times from control trials were not included here as there was no DS or OS. (**C**) Putative DA neurons were identified based on the average waveform of the recorded action potentials. Each dot indicates the amplitude ratio and half width duration of one recorded unit. Two clusters were created using the k-means algorithm to obtain putative DA neurons (blue, n=41) and non-DA neurons (red, n=55).

#### Stage 1: Reward magazine training

Rats were trained in one 30-min session to lick the reward spigot to obtain saccharin solution. Rats were only able to gain access to saccharine reinforcement at a variable interval time schedule in which the first lick after 2, 4, 8 or 16 s (pseudorandomly chosen on each trial) was reinforced. A lick made after one of the four variable time schedules was simultaneously followed after 2 msec by the presentation of a conditioned stimulus, the onset of the reward magazine light (RL). This was followed by the delivery of 0.05 mL (0.05 mL ⁄ s) of sodium saccharin solution (0.25% w ⁄ v) whilst the RL was continuously presented.

#### Stage 2: Modified FR1 training

Rats were then introduced to bar pressing for reward delivery on a modified FR1 (fixed-ratio responding) schedule of lever pressing for 60 minutes. Reward delivery occurred as in the previous stage (including that the RL continued to signal reward), except for two changes to a standard FR1. First, rats were able to gain access to reward by licking on a variable interval time schedule of 32s and 64s (randomly chosen on each trial), to keep the rats active and exploring. Second, a lever was protruded for the entire 60 minutes, and each bar press was followed by the delivery of sodium saccharin solution. Thus, rats were able to gain reward either through licking during the variable time schedule or by bar pressing (at any time). Rats that completed 50 trials by either bar pressing or licking (16 out of 16 rats) moved on to the next stage of training.

#### Stage 3: Standard FR1 training

Reward delivery did not occur by licking of the reward spigot as in stages 1 and 2. Rats were only rewarded when bar pressing for reward on a FR1 schedule (1 bar press=reward delivery) over 60 minutes. An arbitrary maximum of 50 reinforced bar presses in one session was introduced, after which the rats were advanced to the next stage of training. Rats that did not achieve 50 responses (2 out of 16 rats) were given further sessions until they reached the criterion.

#### Stage 4: Discriminative stimulus training

The next stage was designed to place lever pressing under the control of a DS. This stage was conducted over the next ~7 sessions. The rats were split pseudorandomly into two groups, either being presented with a tone or houselight as a DS indicating to the rat the active contingency between bar-pressing and reinforcement. Each session lasted 60 minutes, with 30s blocks with DS on (indicating bar-pressing would lead to reinforcement) pseudorandomly interleaved with 30s blocks with the DS off (indicating bar-pressing would not be reinforced). Rats were hence only rewarded when bar pressing under the 30s continuous presentation of the DS.

Bar presses under DS presentation *versus* no DS were recorded as the discrimination index (bar presses under DS/(bar presses under DS + bar presses under no DS)). Rats (16 out of 16) that achieved > 80% discrimination index were advanced to the next stage.

#### Stage 5: Occasion setting training

The final stage of training was conducted over the next 11 sessions. There were four types of trials in each 60-min session. Only one type of trial allowed the rat access to the sodium saccharin solution. The four types of trials consisted of the following and were presented pseudorandomly (see *Figure 1*): (1) the DS for 30s with no reinforcement of bar pressing; (2) the OS for 30s with no reinforcement of bar pressing; (3) neither the OS nor the DS for 30s with no reinforcement of bar pressing; and (4) the OS for 10s followed by OS+DS for 30s during which time bar pressing was reinforced. The choice of a long interval between the OS and DS (and hence reward opportunity) was motivated by previous research which demonstrated that longer intervals between stimuli (and different sensory modalities; e.g. auditory and visual) favour the acquisition of occasion setting (Fraser and Holland, 2019).

### 2.5. Surgery

Following behavioural training, rats underwent surgery to implant an electrode array that was affixed onto the skull. Rats were anaesthetized using mixture of Isoflurane (5% for induction, 2% for maintenance) and oxygen (1.0 L ⁄ min). A presurgical nonsteroidal, nonopiate analgesic Rimadyl™ (0.5 mL ⁄ kg; 5% w⁄v carpofen; Pfizer Ltd, Kent, UK) was injected subcutaneously. In order to lower the electrode array into place, a hole was drilled stereotaxically at the top of the ventral tegmental area (VTA; 5.80 mm posterior and 0.8 mm lateral to bregma; 7.4–8.4 mm ventral to skull surface).

In addition, five to seven holes were drilled around the area to which the electrode array would be attached, tapped for retaining screws (0–80 hex head, cup point set screws, 1 ⁄ 4 inch; Small Parts Inc.). Using the stereotaxic arm, the electrode array was lowered and dental acrylic used to secure the array to the cranium.

### 2.6. Histology

The following procedure is based upon previous work (Wilson and Bowman, 2006). Rats underwent ~3 weeks of neurophysiological recording, and were killed by an overdose of 0.8 mL Dolethal TM (200 mg ⁄ L pentobarbitone sodium BP; Univet Ltd, Oxford, UK). Following death, they were perfused intracardially with 0.1% phosphate-buffered saline, plus a fixative (4% paraformaldehyde in 0.1 m phosphate buffer).

A freezing microtome was used to cut sections 50 µm thick. These sections were then collected in 0.1 m phosphate buffer, and every fourth was stained for tyrosine hydroxylase and Nissl bodies using standard protocols. In order to conform the position of the electrode microwires with reference to the VTA, all stained sections were analysed under a light microscope and mapped onto standardized sections of the brain (Paxinos and Watson, 2006). A reconstruction of the position of the electrodes is shown in Figure 1B.

### 2.7. Data analysis

To identify responses in the population of putative DA neurons, we compared firing rates across different trial types and time points (Figures 3 and 4). After aligning the activity of each unit to the respective task event (e.g. OS, DS, or bar press), we determined the mean activity of each unit. These mean firing rates were then obtained at different time points around the event of interest, ranging from −5s to +5s relative to the event using 200ms wide window moving in steps of 50ms. At each time point the distributions of firing rates were then statistically compared across conditions using a two-sided Wilcoxon signed rank test (signrank.m function in Matlab). To correct for multiple testing, we applied a p-value threshold that yielded a false discovery rate of 0.05 (Benjamini and Hochberg, 1995).

**Figure 3:**
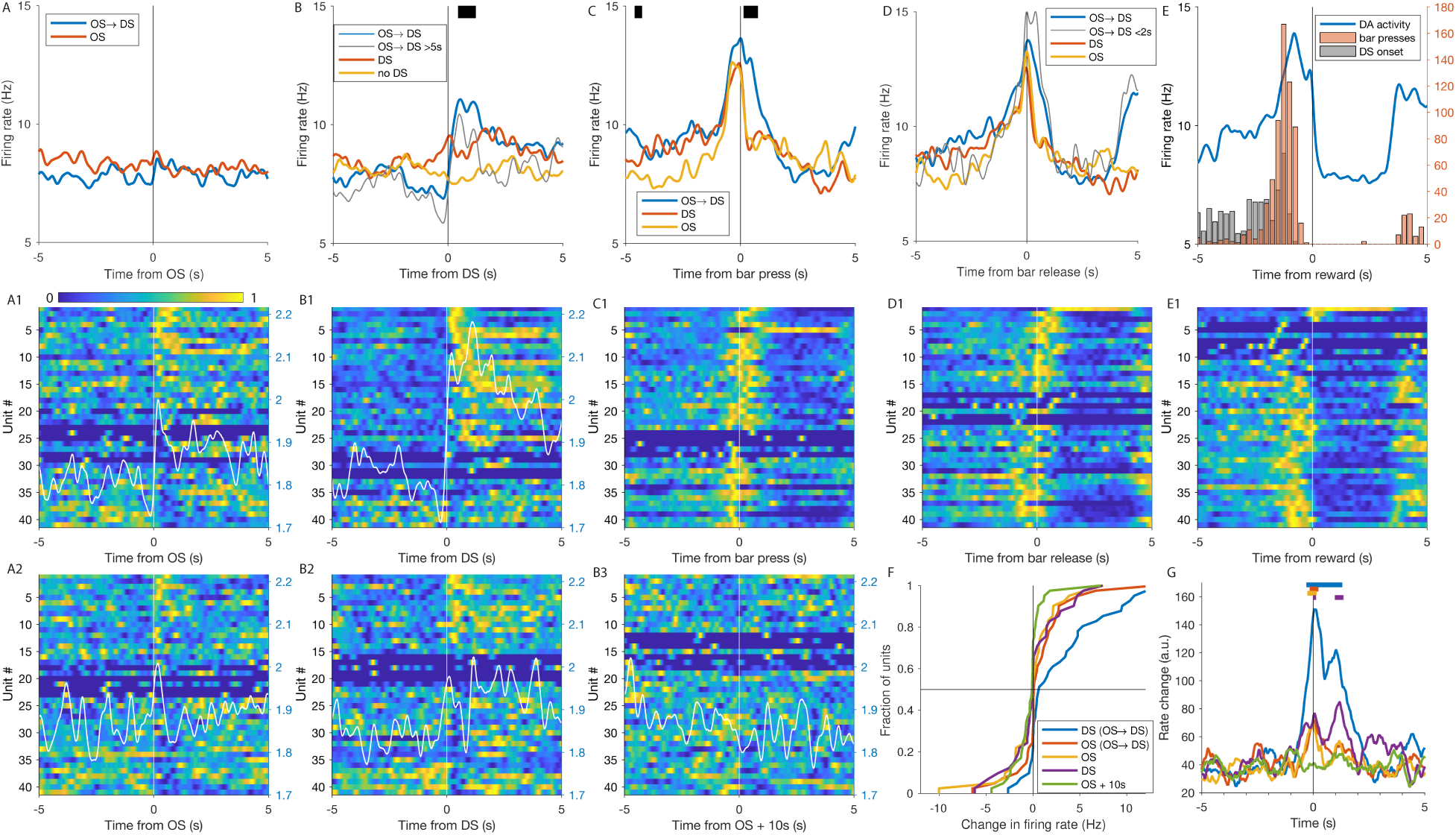
Activity of putative DA neurons in the task. (**A**) Mean firing rate of the population of putative DA neurons around the OS in OS→DS and OS trials. (**A1, A2**) Corresponding activity of each individual putative DA neuron is shown for OS→DS (A1) and OS (A2) trials. Activity of each unit has been normalized for visualisation between 0 (minimal firing rate) and 1 (maximal firing rate; see colorbar). Unit order has been sorted according to the change in activity in the 1s time window preceding the stimulus to the 1s time window following the stimulus (largest increase at the top; units sorted in each panel independently). White line shows the mean of the absolute value of the z-score across all units (right-side scale) (**B**) Mean firing rate of the population of putative DA neurons around the DS in OS→DS and DS trials. The thin grey line shows the activity in the subset of OS→DS trials in which the response time was longer than 5s, i.e. the DS onset and first bar press are separated by more than 5s. The ‘No DS’ trials were the same as OS trials, but here we aligned activity to the time point when the DS would have been presented if this were an OS→DS trial. (**B1-B3**) Corresponding activity of each individual putative DA neuron is shown for OS→DS (B1), DS (B2), and ‘no DS’ control (B3) trials (same visualisations as in A1 and A2). (**C**) Mean firing rate of the population of putative DA neurons around the first bar press following the DS in OS→DS trials, following the DS in DS trials, and following the OS in OS trials. (**D**) Mean firing rate of the population of putative DA neurons around the first bar release following the DS in OS→DS trials, the DS in DS trials, and the OS in OS trials. The thin grey line shows activity in the subset of trials in which the preceding DS occurred more than 2s ago. In (A-D) Black bars at the top indicate time points when there was a significant difference between activity in the OS→DS and OS trials (A) or between the OS→DS and DS trials (B-D) (two-sided Wilcoxon signed rank test; 200ms time windows with a 50ms step size, see Methods). (**E**) Mean firing rate of the population of putative DA neurons around reward. Overlaid histogram indicates the number of bar presses and DS onset relative to the reward (event counts given on the right side y-axis). (**C1-E1**) Each panel shows the corresponding single unit activity for bar press, release and reward events, respectively. (**F**) Firing rate changes from 1s before to 1s after the OS or DS are illustrated using their cumulative distributions. (**G**) Absolute rate changes summed over all units at different time points verify that at t=0 (i.e. OS or DS onset) there is significant change in activity (same legend as in panel F). Colour bars at the top indicate time points when the rate change was significant with respect to a permutation test using the 10s preceding the stimulus as a baseline (see Methods for details).

**Figure 4:**
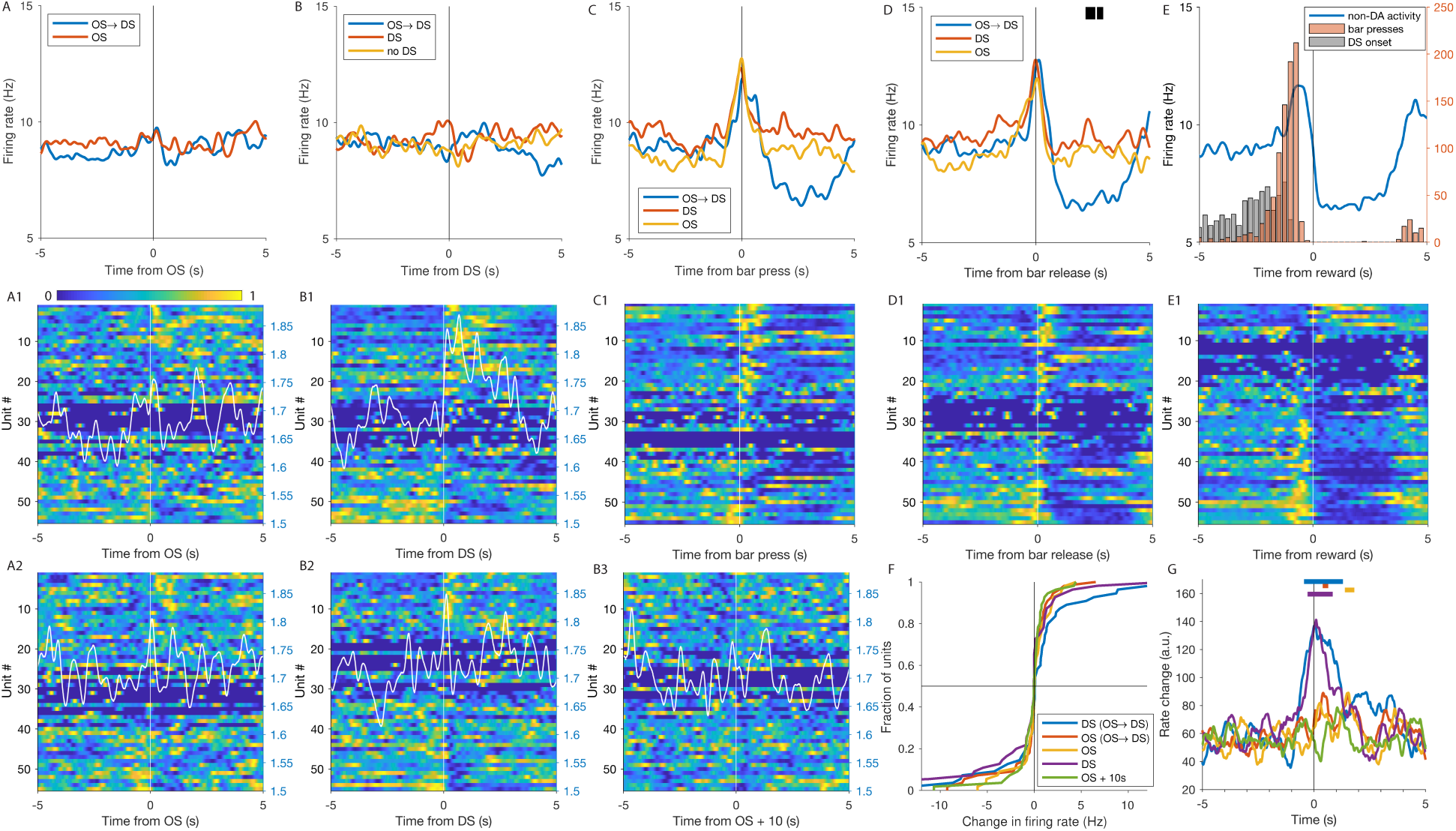
Activity of putative non-DA neurons in the OS paradigm. (**A-E**) Mean firing rate of the population of putative non-DA neurons in the different trial types and task events (same analyses as in Figure 3). (**A1-E1**) Corresponding activity of each individual putative non-DA neuron is shown for all trial types and events (same analyses as in Figure 3). (**F**) Firing rate changes from 1s before to 1s after the OS or DS are illustrated using their cumulative distributions. (**G**) Absolute rate changes summed over all units at different time points verify that at t=0 (ie OS or DS onset) there is significant change in activity (same legend as in panel F). Colour bars at the top indicate time points when the rate change was significant with respect to a permutation test using the 10s preceding the stimulus as a baseline (see Methods for details).

To visualize the mean firing rate of all individual units (Figures 2 and 3) we normalized their activity to a range between 0 and 1 by first subtracting their minimal firing rate and then dividing by the maximal firing rate of the result. The minimal firing rate was obtained within the 10s time window around the shown event.

For plotting, the individual and the population mean firing rates were smoothed using a 500ms wide Gaussian window with a standard deviation of 100ms. Statistical tests were performed on the firing rates before smoothing. For the visualization of the mean absolute values of the z-score of the firing rates (Figures 3 and 4), the session-wide mean and standard deviation of the firing rate of each unit was determined in 1s time windows and then used to calculate the z-score.

For the permutation analysis in Figures 3G and 4G we aimed to determine the distribution of rate changes, which occur randomly. To do this we calculated rate changes at random time points relative to the event of interest. Each time point was drawn from a uniform distribution covering 9s before to 9s after the event. For a given permutation a time point was drawn for each unit individually. The rate change was then calculated as the mean of the firing rate across the 1s time window after each time point minus the mean firing rate across the 1s time window preceding time point. For each iteration this procedure yielded a distribution of rate changes, with one value for each unit. We then summed up the absolute values of these rate changes and repeated this procedure 10,000 times for each trial type and stimulus alignment of interest (i.e. DS in OS→DS trials, OS in OS→DS trials, OS in OS trials, DS in DS trials, and 10s after the OS in OS trials). This approximated a distribution of absolute rate changes that are expected just by chance. We then compared the actual absolute rate changes (shown in Figures 3G and 4G) with the corresponding distribution obtained from the permutation procedure. At each time point this yielded a p-value, given by the fraction of permutated rate changes that were equal or larger than the empirical ones. Here we used a simple Bonferroni correction based on five different trial types / stimuli and ten independent 1s time windows for the rate change calculation, yielding a p-value threshold for significance of 0.001.

### 2.8. Neurophysiology

#### Spike sorting

Spikes were firstly sorted online in Spike 2 ™ version 6 (Cambridge Electronic Design, Cambridge, UK) using the waveform shape template matching, and re-sorted offline by performing principal components analysis to visualise waveform clusters. The first three principal components of each spike were assigned a co-ordinate in 3-D space, to be able to cluster similar waveforms together. Separate clusters were then classified using the Normal Mixtures algorithm in Spike2 (modified to include waveforms 2.5-3 standard deviations of the centroid of that cluster as indicated by the best discrimination among clusters as judged by visual inspection). Finally, overlaid waveforms were visually inspected to reject any putative spike that seemed to be the result of a mechanical or electrical noise. The quality of the clustering was assessed by calculating the signal-to-noise ratio within each cluster. Single neurons were classified using the following criteria: there were no signs of noise at 50 Hz or its harmonics, the inter-spike interval histogram exhibited a refractory period, and there were no electrical artefacts within the cluster from the rat bar pressing or licking the spigot.

## 3. Results

### 3.1. Behaviour

The behavioural task consisted of four different trial types (Figure 1A). The first trial type (“OS→DS”) started with the onset of the OS, followed after 10s by the onset of the DS. Once the DS was presented, any bar press activated the reward magazine and delivered the reward to the animal. Bar presses before the DS had no effect. The second trial type (“OS”) started in the same way, with the onset of the OS. However, there was no subsequent DS, and bar presses had no effect. In the third trial type (“DS”) there was no OS, but instead it started immediately with the DS, and bar presses were again not rewarded. Finally, some trials were control trials (“none”), without any stimuli (i.e. no OS or DS) or rewards.

To see whether the behaviour of the animals reflected learning of the task, we determined the number of bar presses per minute in the different trial types. We found that in OS→DS trials, after the onset of the DS, the rate of bar presses was significantly higher compared to all other trial types (p=2.4*10^−5^ for OS trials; p=2.8*10^−7^ for DS trials; p=4.1*10^−20^ for ‘none’ trials, one-tailed 2-sample Kolmogorov-Smirnov tests; Figure 2A). This indicated that the animals had learned the overall task, and in particular that bar presses were only rewarded when the DS was preceded by the OS. Furthermore, we analysed response times, i.e. the time it took the animals to respond to the onset of a stimulus with a bar press. Learning would be reflected in fast response times to the DS in OS→DS trials, compared to OS and DS control trials, respectively. Indeed, we found that OS→DS trials had significantly faster response times (median=3.85s, MAD=2.26s) than OS and DS trials (p=1.6*10^−6^ for OS trials, median 4.6s, MAD=3.0s; p=2.5*10^−7^ for DS trials, median=3.75s, MAD=2.3s, one-tailed 2-sample Kolmogorov-Smirnov tests; Figure 2B). Based on these results we conclude that the animals used the OS and DS to guide their behaviour as intended by the task design.

### 3.2. Cell type classification

Neural activity was recorded in the VTA as verified by histological analysis of the electrode tracks (Figure 1B), and then spike sorted based on the waveform. To determine whether recorded units corresponded to putative DA neurons, we analysed the shape of the average waveform of each recorded unit. Using the amplitude ratio and half width duration to cluster the units (see Methods) yielded a distinct profile of putative DA (n=41) and non-DA neurons (n=55; Figure 2C), similar to previous studies (Pan, Schmidt, Wickens, and Hyland, 2008; Roesch, Calu, and Schoenbaum, 2007).

### 3.3. Neural Responses to the Occasion Setter and Discriminating Stimulus

We investigated whether putative DA neurons distinguished the reward-predicting properties of the OS and DS in the different trial types.

Firstly, we aligned the activity of each putative DA neuron to the onset of the OS, which occurred in two trial types (OS→DS and OS). In OS→DS trials, 10s after the onset of the OS, the DS was presented. In contrast, in OS trials, the DS never occurred. The onset of the OS did not elicit a visible response in the mean firing rate of the population of putative DA neurons in either trial type (Figure 3A). However, inspection of the individual unit firing rates (Figure 3, A1 and A2) indicated the presence of both increases and decreases as a response to the OS. We visualized these apparent responses using the absolute value of the mean z-score as a measure of activity changes, so that increases and decreases in firing rate would summate instead of cancelling each other out (shown as white lines in Figure 3, A1 and A2). As the amplitude, timing and duration of these responses varied considerably across units, we employed a more elaborate analysis instead of attempting to count the number of units with significant increases and decreases (see below). We conclude that despite the OS being a behavioural significant event that the animals used to guide their behaviour, there was no population increase in putative DA cell activity. However, individual neurons responded to the OS, potentially reflecting an intermediate stage of learning, or a more local mode of DA signalling.

Secondly, we aligned activity of each putative DA neuron to the onset of the DS. Again, this could be done for two trial types, OS→DS trials and DS trials (i.e. with no preceding OS). For OS→DS trials there was a sharp response to the onset of the DS with a ~4Hz increase in the mean firing rate of putative DA neurons (Figure 3B). This increase was significant compared to the mean firing rate in DS trials (Figure 3B; two-sided Wilcoxon signed rank test, see Methods), and compared to the activity 10s after the onset of the OS in OS trials (‘no DS’ in Figure 3B). The response to the DS was also present in the subset of OS→DS trials, in which it took the animals more than 5s to press the bar afterwards (Figure 3B). Therefore, the sharp increase in OS→DS trials was not due to subsequent bar presses, but a response to the DS itself.

Examining the corresponding single-unit activity patterns revealed a strong response in OS→DS trials in the majority of units (Figure 3, B1). While in DS trials there was no prominent increase in the mean firing rate, individual neurons showed increases or decreases (Figure 3, B2), very similar to the OS responses described above. This was supported by the corresponding visualisation of the absolute value of the mean z-score, yielding a sharp increase at DS onset. As a control we used again the ‘no DS’ trials and examined activity 10s after the onset of the OS, the time when the DS would have occurred. Importantly, in this control there were no marked increases or decreases in the firing rate of individual neurons (Figure 3, B3), also supported by the absence of an increase in the absolute mean z-score. These results demonstrate that the response to the DS can be flexibly modulated by the OS to either yield a population increase or a population ‘null’ signal.

We then looked into more detail into the single unit responses to the OS and DS. Note that also for simple baseline activity, random fluctuations would lead to a spurious response pattern when the units are sorted based on their activity (Figure 3). Therefore, we devised additional analyses to ensure that the observed single-unit increases and decreases exceed chance level. First, for each unit, we determined the change in firing rate in the 1s time window preceding the stimulus to the 1s time window following the stimulus. The resulting distribution of firing rate changes (one value per unit) was then compared across the different stimuli and trial types, using their cumulative distributions for visualisation (Figure 3F). For OS→DS trials, there was a strong response in the majority of individual units (Figure 3, B1), which was reflected in the cumulative distribution of the firing rate changes being shifted towards positive values (Figure 3F). At the other extreme, our ‘no DS’ control condition (10s after the OS onset), with no apparent single-unit responses (Figure 3, B3), yielded a more symmetrical and narrow cumulative distribution of firing rate changes (Figure 3F). In contrast the responses to the OS (in OS→DS and OS trials) and DS (in DS trials) were wider, reflecting the presence of units with larger firing rate changes, both positive and negative ones. Next, we calculated the sum of all absolute firing rate changes across all units in a given condition, obtaining a measure for population rate changes that is sensitive to parallel increases and decreases in firing rate. This measure was calculated at different time points relative to the onset of the cue (OS or DS). The resulting modulations over time showed that there was a significant change in firing rate at the onset of the OS and DS (permutation test, see Methods; Figure 3G). In contrast, for our ‘no DS’ control, there was no such change. This verifies our assertion above that, despite the absence of increases in mean firing rate, the OS (in OS→DS and OS trials) and DS (in DS trials) lead to increases and decreases in the firing rate of individual DA neurons.

### 3.4. Bar pressing and reward responses

Next we analysed neural responses related to the bar pressing behaviour of the animals. We examined activity related to bar presses in three trial types: OS→DS, DS, and OS trials. To reduce the impact of variability across bar presses in terms of timing and reward predictions, we focussed on the first bar press that the animals performed after the onset of the DS or OS. We found that in OS→DS trials, putative DA neurons increased their firing rate around bar presses, starting about 1s before the bar press, peaking at the time of the bar press, and lasting until about 1s after the bar press (Figure 3C). Unrewarded bar presses in OS or DS trials had a similar temporal profile preceding the bar press, but after the bar press, the firing rate dropped quickly back to baseline, yielding significantly different firing rates up to 1s after the bar press (Figure 3C). This might be due to the lack of reward feedback in the DS and OS trial types compared to the OS→DS trials.

Although the animals typically released the bar quickly after the bar press, there was some temporal variation between these events. Aligning activity to the bar release, yielded a similar temporal profile in the firing rate increase across trial types (Figure 3D). As the bar release immediately triggered the reward magazine light, it could be considered a key event in the behavioural sequence leading to the reward. Seemingly, the temporal profile of the putative DA neurons corresponded to a slow ramping increase in firing rate starting up to 5s before the bar release (Figure 3D). As this ramping increase in firing rate was reminiscent of the increases in striatal DA concentration occurring over seconds during goal approach (Howe, Tierney, Sandberg, Phillips, and Graybiel, 2013) and reinforcement learning (Hamid, Pettibone, Mabrouk, Hetrick, Schmidt, Vander Weele, Kennedy, Aragona, and Berke, 2016; Mohebi et al., 2019), we examined it in further detail. Our results indicated the ramping firing rate in this case was due to the averaging across trials with different event timings, rather than a slowly ramping, putative motivational signal. This was supported by a control analysis on the subset of trials in which the DS onset occurred within less than 2s before the bar release. In this control analysis the apparent ramp occurred only within the 2s before the bar release (Figure 3D), supporting that the ramp is merely due to preceding DS and bar press events (Figure 3B, C). Similarly, the activity, when aligned to the onset of the reward, exhibited a ramp over seconds that basically matched the distribution of preceding bar presses and DS onsets (Figure 3E). In contrast to the OS and DS responses described above, the single-unit firing rates related to the bar press, bar release, and reward were very homogeneous across the population of putative DA neurons (Figure 3, C1-E1).

We conclude that also in this operant paradigm, involving contextual OS cues and longer time scales, DA cell firing rates exhibited brief firing rate increases to reward-predictive cues and actions, but did not show ramping activity as presumed for motivational signals.

### 3.5. Comparison with non-DA neurons

We repeated all analyses of the firing rates in the different trial types for neurons that were labelled as putative non-DA neurons (Figure 2C). Similar to the putative DA neurons, also the non-DA units exhibited no clear response to the OS in the population mean firing rate (Figure 4A). However, also the inspection of the individual unit activity and the mean absolute z-score (Figure 4, A1 and A2) did not suggest the existence of any responses on a single unit level.

In contrast to the DA analyses, the non-DA units did not show any population response to the DS (Figure 4B). However, interestingly, on a single unit level DS responses were present, which seemed to cancel out in the population average. For the OS→DS trials these responses occurred mostly within 3s after the DS, while in DS trials the responses seemed to occur only within less than a second after the DS (Figure 4, B1 and B2). The responses to the bar press and release had a similar time course as the DA units, but showed no significant differences between the trial types (Figure 4C, D). This seemed to be mostly due to overall briefer responses, which were also visible on a single unit level (Figure 4, C1 and D2). Finally the reward responses exhibited a similar profile as the DA units, but the non-DA units decreased their activity briefly before the reward onset (Figure 4E, E1), while the DA units decrease occurred a bit later.

For the analysis of the firing rate changes from −1s to +1s around the stimulus, the cumulative distributions of the rate changes strongly overlapped across the different conditions (Figure 4F). However, the statistical analyses of the rate changes revealed a significant rate change around the time of the DS in both OS→DS and DS trials (Figure 4G). In contrast, rate changes in OS→DS and OS trials did not show pronounced increases at OS onset, indicating that they did not elicit any or only weak/short responses to the OS.

We conclude that putative non-DA neurons did not show population level responses to reward-predicting stimuli in the OS paradigm. However, on a single-unit level they did show both increases and decreases to the DS, which cancelled out on a population level. The absence of OS responses in putative non-DA neurons suggests that the single-unit responses to the OS in putative DA neurons originate from external inputs rather than from other VTA neurons.

## 4. Discussion

Our results showed that putative DA neurons encoded the relationship between the OS and the DS. Responses of putative DA cells to the DS were enhanced when preceded by the OS, as were behavioural responses to obtain rewards. This is consistent with the hypothesis that responses of DA neurons to the DS contribute to the expression of behavioural responses. Surprisingly though, we did not find a population response of putative DA neurons to the OS, contrary to predictions of standard temporal-difference models of DA neurons. As the OS was the earliest predictor of reward in the task sequence of events, standard temporal-difference models would predict at least a partial shift of the DA response from the reward to the OS (Pan et al., 2005; Schultz et al., 1997). Intriguingly, despite the absence of a population response, our recorded putative DA neurons exhibited a heterogeneous response on a single unit level, so that some units increased and others decreased their activity as a response to the OS. Similarly, putative non-DA cells did not respond to the DS on a population level, but with heterogeneous responses on a single unit level.

DA cell responses have not been studied so far in tasks with OSs. Our behavioural task closely followed the pivotal studies that introduced serial feature positive discrimination (Looney and Griffin, 1978; Ross and Holland, 1981; Sainsbury and Jenkins, 1967). In their procedure the OS is presented before the DS, which is then followed by a behavioural response leading to reward. However, when the DS was presented on its own, the response is not rewarded. The behavioural analysis in these classic studies revealed that conditioned responding to the OS was minimal compared to the DS when preceded by the OS. This demonstrated that the OS, rather than creating a simple associative link with reward, modulated the response determined by the DS. Our task and behavioural data matched these findings, permitting us to address the underlying neural processes involved DA signalling.

Our neurophysiological data indicated that midbrain DA cells might act as neurobiological substrate for encoding occasion setting properties, perhaps in coordination with neurons in the OFC (Shobe, Bakhurin, Claar, and Masmanidis, 2017). Furthermore, our results pointed to a limitation of standard temporal-difference models to account for DA cell responses to reward predicting stimuli, and support models that employ a more complex state representation (Daw, Courville, and Touretzky, 2006). There are several possibilities for the lack of neural responses to the OS at a population level. Firstly, in our task there was a long time delay between the OS and the reward. In classical conditioning DA cells of primates reduce their responses to the conditioned stimulus hyperbolically as a function of the interval between the conditioned and unconditioned stimulus (Kobayashi and Schultz, 2008). However, albeit in that study DA responses were reduced for long intervals, DA neurons still fired to conditioned stimuli that were 16s away from the reward, which would be within the time range of the OS-reward interval in our paradigm. Another difference was that in Kobayashi and Schultz (2008) DA neurons increased their firing rate following reward presentation, a pattern that we did not observe in our data. Instead during the presentation and consumption of the reward, our putative DA cell activity decreased (Kiyatkin and Gratton, 1994; Nishino, Ono, Muramoto, Fukuda, and Sasaki, 1987; Richardson and Gratton, 1996). Secondly, the lack of the population response to the OS could be due to the low contingency between the OS and the reward. The OS predicted the reward with only 50% probability, and was necessary but not sufficient for reinforcement. Responses of DA neurons to conditioned stimuli encode reward probability, with stronger responses for higher reward probabilities (Fiorillo et al., 2003). However, this would only explain a weak response to the OS, rather than the absence of a population response. Thirdly, although DA neurons did not respond to the OS at a population level, they did respond with increased and decreased firing in the activity of single units. This suggests that context, here in the form of the OS stimulus, may be encoded by DA neurons at a single-cell level. Although this contrasts with the idea of DA cells providing a global signal to the striatum, this type of signalling may be relevant for more localized changes in striatal DA concentration. To determine this future studies could examine whether such heterogeneity in the DA cell responses (Fiorillo, Yun, and Song, 2013) is reflected in different anatomical subgroups and projection targets, which we could not assess in the present data.

DA cell activity is often characterized by brief phasic increases. While slow, ramp-like increases in DA concentration have been found in the striatum (Hamid et al., 2016; Howe et al., 2013), corresponding slow increases in the firing rate of DA neurons have not been found (Mohebi et al., 2019). One potential explanation for this discrepancy is that DA cell firing has typically been studied in simple behavioural tasks such as classical conditioning (Pan et al., 2005), while striatal DA concentration has also been measured in more complex tasks (Hamid et al., 2016), involving longer time scales for approaching a goal (Howe et al., 2013). In the present study longer timescales (10s of seconds) were required for the animal to integrate information about the OS and DS, and the reward contingencies were more complex than in a classical conditioning paradigm. Still, our analyses on the firing rates suggested that DA cell responses consisted mostly of phasic changes in relation to the OS, DS, and bar pressing. Therefore, we conclude that also in this more complex task involving longer time scales, DA cell firing showed no evidence for ramp-like increases in firing rate that might correspond to a motivational signal.

In summary, DA neurons incorporated context in their responses to reward-predicting stimuli and exhibited complex responses to OS stimuli that point to heterogeneous rather than global DA signals.

## Acknowledgments

This research was supported by a doctoral studentship to Luca Aquili from the UK Engineering and Physical Research Council.

## Abbreviations

DA: Dopamine
DS: Discriminative stimulus
MAD: Median absolute deviation
OS: Occasion setter
VTA: Ventral tegmental area

